# Is a large eye size a risk factor for myopia? A Mendelian randomization study

**DOI:** 10.1101/240283

**Authors:** The UK Biobank Eye and Vision Consortium.

## Abstract

Myopia (nearsightedness) is an increasingly common cause of irreversible visual impairment. The ocular structures with greatest impact on refractive error are corneal curvature and axial length. Emmetropic eyes range in size within and across species, yet possess a balance between corneal curvature and axial length that is under genetic control. This scaling goes awry in myopia: 1 mm axial elongation is associated with ~3 Dioptres (D) myopia. Evidence that eye size prior to onset is a risk factor for myopia is conflicting. We applied Mendelian randomisation to test for a causal effect of eye size on refractive error. Genetic variants associated with corneal curvature identified in emmetropic eyes (22,180 individuals) were used as instrumental variables and tested for association with refractive error (139,697 individuals). A genetic risk score for the variants was tested for association with corneal curvature and axial length in an independent sample (315 emmetropes). The genetic risk score explained 2.3% (P=0.007) and 2.7% (P=0.002) of the variance in corneal curvature and axial length, respectively, in the independent sample, confirming these variants are predictive of eye size in emmetropes. The estimated causal effect of eye size on refractive error was + 1.41 D (95% CI. 0.65 to 2.16) *less* myopic refractive error per mm flatter cornea (P<0.001), corresponding to +0.48 D (95% CI. 0.22 to 0.73) more hypermetropic refractive error for an eye with a 1mm longer axial length. These results do not support the hypothesis that a larger eye size is a risk factor for myopia. We conclude the genetic determinants of normal eye size are not shared with those influencing susceptibility to myopia.

## Introduction

Myopia (nearsightedness) occurs when the eye focuses light from distance objects in front of the retina, resulting in an inability to obtain a clear image of objects far away. A characteristic feature of myopic eyes is that the combined refractive power of the cornea and crystalline lens is too high in relation to the axial eye length; in most cases the cause is an excessively elongated eye [1]. The prevalence of myopia has increased dramatically in recent decades, especially in parts of East and Southeast Asia [2, 3]. This has important public health implications, since myopic eyes are at greater risk of retinal detachment, choroioretinal atrophy, glaucoma and certain types of cataract, which together make it a leading cause of visual impairment and blindness [4, 5].

Two important environmental risk factors for myopia have been identified to date – education and (insufficient) time spent outdoors in childhood [6–9] – and more than a hundred genetic loci that influence susceptibility to myopia have also been discovered [10–12]. Despite this progress, little is understood about the mechanisms linking genetic variants and environmental exposures to the excessive elongation that upsets the usual balance and scaling of the eye's component parts.

One line of enquiry has reasoned that the cellular and molecular pathways responsible for determining normal eye size are invoked to increase axial length in myopia. In support of this theory, a genetic correlation has been observed between axial length and refractive error [13, 14], implying that a shared set of genetic variants plays a role in determining both traits. Furthermore in some studies, infants and children destined to become myopic have been found to have longer eyes even before myopia develops, i.e. eye length has been shown to be predictive of myopia development [15, 16]. However, arguing against this theory, axial length was not predictive of myopia development in a further study [17], and in a sample of chicks with experimentally-induced myopia, the genetic correlation between pre-treatment eye size and myopia susceptibility was very close to zero [18], suggesting that different sets of genetic variants control myopia and normal eye size.

Mendelian randomisation is a powerful approach for estimating the causal effect of an exposure on the risk of a disease or other outcome. The approach exploits genetic variants robustly associated with an exposure as instrumental variables for assessing an exposure-outcome relationship; unlike conventional ("observational") estimates of exposure-outcome relationships, causal estimates from Mendelian randomisation analysis are free from bias due to reverse causation and less susceptible to bias from unmeasured confounders [19, 20].

Here, in order to gain insight into the related questions (1) is eye size in childhood predictive of myopia development, and (2) are the molecular pathways that normally regulate eye size also used to produce an enlarged myopic eye, we used a Mendelian randomisation framework to test the hypothesis that genetic variants responsible for controlling the normal variation in eye size in emmetropes also cause susceptibility to myopia.

## Methods

### Study cohorts and genotype data quality control

#### UK Biobank

The UK Biobank is a longitudinal study of the health and well-being of approximately half a million UK residents [21]. Ethical approval was obtained from the National Health Service (NHS) National Research Ethics committee (Ref. 11/NW/0382) and all participants provided informed consent. Participants were recruited between 2006–2010, when they attended 1 of 22 assessment centres distributed across the UK, and completed a series of interviews and physical or cognitive measurements. Approximately 25% of participants underwent an ophthalmic assessment, which was introduced towards the latter stages of recruitment. This included a logMAR visual acuity (VA) examination at a test distance of 4 metres, with habitual spectacles if worn, and non-cycloplegic autorefraction/ keratometry (Tomey RC5000; Tomey GmbH Europe, Erlangen-Tennenlohe, Germany).

Participants were excluded from the analyses if they had a history of an eye disorder that may have altered their physiological refractive error or corneal curvature. Specifically, individuals were excluded if they self-reported a history of laser refractive surgery, cataract surgery, corneal graft surgery, any other eye surgery in the last 4 weeks, any eye trauma resulting in sight loss, serious eye problems, or self-report of having cataracts or retinal detachment. Participants were also excluded if their hospital records indicated they had undergone cataract surgery, retinal detachment surgery, or corneal surgery.

UK Biobank researchers extracted DNA samples from blood, genotyped the samples on either the UK BiLEVE array (n=49,950) or the UK Biobank Axiom array (n=438,427) and imputed to the HRC reference panel and a combined 1000 Genomes Project-UK10K reference panel using IMPUTE4 [22]. Imputed genotype data were available for 488,377 participants (June 2017 release; see Bycroft et al. [22]). We classified individuals as having European vs. non-European ancestry using the results of principal components (PC) analysis. First, a set of unrelated individuals from the n=409,728 White British ancestry subset defined by Bycroft et al. [22] were filtered to exclude heterozygosity outliers (autosomal heterozygosity more than ±4 standard deviations (SD) from the mean level). Next, we calculated the mean and SD for each of the top 20 PCs in this sample of unrelated White British ancestry individuals. Finally, we defined as European all individuals who fell within the mean ±10 SD for each of these top 20 PCs [23] and who also self-reported their ethnicity as White, British, Irish or any other white background. This resulted in a total of 443,400 individuals meeting our criterion of European ancestry, some of whom were related.

#### CREAM Consortium

The CREAM Consortium carried out a meta-analysis of refractive error GWAS studies [24]. All participants provided informed consent during recruitment into the individual studies [24]. Here, we restricted attention to GWAS studies carried out in participants of European ancestry using the Spherical Equivalent phenotype, measured in Dioptres. All participants were aged >25 years. The combined sample size was n=44,192. All studies imputed genotype data to the 1000-Genomes Project phase 3 reference panel; however not all samples included in the meta-analysis had imputed genotype information for all markers, due to some markers being excluded during per-cohort quality control procedures.

#### ALSPAC (Avon Longitudinal Study of Parents and Children)

Pregnant women resident in Avon, UK with expected dates of delivery 01/04/1991 to 31/12/1992 were recruited into the study. Of 14,541 initial pregnancies, 13,988 children were alive at 1 year of age. When the oldest children were approximately 7 years of age, an attempt was made to bolster the initial sample with eligible cases who had failed to join the study originally. This resulted in an additional 713 children joining the study. Ethical approval for the study was obtained from the ALSPAC Ethics and Law Committee and the Local Research Ethics Committees. Boyd et al. [25] have published a profile of the cohort, and the study website contains details of all the data that is available through a fully searchable data dictionary (www.bris.ac.uk/alspac/researchers/data-access/data-dictionary).

As described [26], ALSPAC children were genotyped using the Illumina HumanHap550 quad chip. ALSPAC mothers were genotyped using the Illumina human660W-quad chip. Following quality control (individual call rate >0.97, single nucleotide polymorphism (SNP) call rate >0.95, minor allele frequency (MAF) > 0.01, Hardy-Weinberg equilibrium (HWE) >1.0e-07, cryptic relatedness within mothers and within children identity-by-descent (IBD) <0.1, non-European clustering individuals removed) 8,237 children and 8,196 mothers were retained with 477,482 SNP genotypes in common between them. Haplotypes were estimated on the combined sample using ShapeIT (v2.r644) [27]. Imputation was performed using IMPUTE v2.2.2 [28] against all 2186 reference haplotypes (including non-Europeans) in the Dec 2013 release of the 1000 Genomes Project reference haplotypes (Version 1, Phase 3). Imputed genotype data were available for a total of 8,237 children. Participants who withdrew consent were excluded from our analyses.

ALSPAC participants were invited to attend a number of visits to an assessment centre. The visit held when participants were aged approximately 15 years old included a vision assessment, at which refractive error was measured by non-cycloplegic autorefraction (Canon R50; Canon USA, Inc., Lake Success, NY, USA) and in a subset (the final year of data collection) axial length and corneal curvature were measured by partial coherence interferometry and infra-red keratometry, respectively (IOLmaster; Carl Zeiss Meditec, Welwyn Garden City, UK).

### Selection of instrumental variables for eye size

To identify genetic variants associated with eye size in emmetropes we carried out a GWAS for corneal curvature in emmetropic UK Biobank participants. We defined emmetropic eyes as those with spherical (SPH) and astigmatic (CYL) refractive error of 0.00 ≤ SPH ≤ +1.00 D and 0.00 ≤ |CYL| ≤ +1.00 D, respectively, and with a VA <0.2 logMAR. If both eyes were classified as emmetropic, we took the average corneal curvature of the 2 eyes as the phenotype. If only 1 eye was classified as emmetropic, we took the corneal curvature of that eye as the phenotype. There were a total of 22,180 individuals with at least 1 emmetropic eye who met the criteria for inclusion in the GWAS for corneal curvature; Figure S1 outlines the selection scheme for these participants. Association tests were conducted using BOLT-LMM [29] for 6,961,902 genetic markers present on the HRC reference panel [30] with MAF ≥0.05 and IMPUTE4 INFO metric >0.9 and per-marker and per-individual missing genotype rates <0.02. Age, gender, genotyping array (coded as 0 or 1 for the UK BiLEVE or UK Biobank Axiom, respectively) and the first 10 PCs were included as covariates. The genetic relationship matrix for the BOLT-LMM analysis was created using a set of approximately 800,000 well-imputed variants (INFO >0.9) with MAF >0.005, missing rate ≤0.01, and an 'rs' variant ID prefix that were LD-pruned using the --indep-pairwise 50 5 0.1 command in PLINK 2.0 [31]. The GWAS summary statistics were filtered to remove A/T or G/C variants, markers with a p-value <0.01 for a test of HWE and those not present in the summary statistics from the CREAM consortium refractive error GWAS meta-analysis. A set of independent markers associated with corneal curvature in emmetropes (P<5.0e-08) were selected by sequentially choosing the most strongly-associated marker, excluding all markers within ±500 kb of the top marker or having pairwise linkage disequilibrium (LD) r^2^<0.2 with the top marker, and so on until there were no further markers with P<5.0e-08. This identified 32 markers independently and strongly associated with corneal curvature in emmetropic eyes (Table S2).

### Association of instrumental variables with refractive error

#### Combined CREAM consortium and UK Biobank GWAS results

We carried out a GWAS for refractive error in UK Biobank participants using the methods described above for corneal curvature. We included 95,505 participants of European ancestry who had autorefraction information available and no history of eye disorders (Figure S2). All repeat refractive error readings were averaged after removal of those flagged as unreliable. Mean spherical equivalent (MSE) refractive error was calculated as sphere power plus half the cylinder power. The refractive error of an individual was taken as the average spherical equivalent of the two eyes. BOLT-LMM was used to test for association between refractive error and each of the 6,961,902 genetic markers tested in the corneal curvature GWAS. Age, gender, genotyping array, and the first 10 PCs were included as covariates.

A meta-analysis of the CREAM consortium refractive error GWAS summary statistics (maximum n=44,192) and the above UK Biobank refractive error GWAS summary statistics (n=95,505) was carried out using a fixed effects, standard error-weighted model with the program METAL [32]. Using the meta-analysis results, we obtained the beta coefficient (in units of dioptric change in refractive error per copy of the risk allele) and standard error for each of the 32 markers associated with corneal curvature in emmetropes. All individuals analysed in the corneal curvature GWAS were also included in the UK Biobank refractive error GWAS, hence the degree of sample overlap was 22,180/(95,505 + 44,192) = 16%.

#### CREAM consortium GWAS

For each of the 32 markers associated with corneal curvature (Table S2) we obtained the beta coefficient (in units of dioptric change in refractive error per copy of the risk allele) and standard error from the CREAM GWAS meta-analysis summary statistics. Care was taken to ensure that the risk and reference alleles were matched across the UK Biobank corneal curvature GWAS and the CREAM refractive error GWAS.

### Statistical analyses

Unless otherwise stated, all analyses were carried out using the R statistics program. Inverse variance-weighted, Egger, and median-based Mendelian randomisation analyses were carried out using the MendelianRandomization package (maintained by Olena Yavorska and Stephen Burgess). The variance in corneal curvature or axial length explained by the 32 instrumental variable markers was assessed in ALSPAC participants using ocular data for the children when they were approximately 15 years old. A genetic risk score [33] (also known as an allele score) for the 32 genetic markers was computed for each child using the --score function in PLINK 1.9 [31]. Emmetropic eyes of ALSPAC participants were defined as those with refractive error 0.00 ≤ SPH ≤ +1.00 D and 0.00 ≤ |CYL| ≤ +1.00 D, respectively. Corneal curvature or axial length in emmetropic eyes (averaged between the 2 eyes if both eyes were emmetropic) was regressed on gender in a baseline model. The same phenotype was then regressed on gender plus the polygenic risk score in a full model, and the difference in the adjusted R^2^ between the baseline and full models was calculated. The difference in R^2^ between an analogous full model and a baseline model was also calculated for all participants with available data, i.e. without restriction to emmetropic eyes.

## Results

### Relationship between axial length and corneal curvature in emmetropes vs. non-emmetropes

The relationship between axial length and corneal curvature in emmetropic and non-emmetropic eyes has been reported in several prior studies [34–39]. As an illustration of these relationships, Figure 1 depicts data for 15-year-old participants in the ALSPAC (note that axial length was not assessed in the UK Biobank, hence comparable plots were not available for this larger study cohort). In eyes classified as emmetropic, corneal curvature and axial length exhibited a consistent linear association; the axial length:corneal curvature ratio was 2.943 (95% CI. 2.935 to 2.952; n=306). By definition, eye size and refractive error were only weakly associated in these emmetropic eyes (Figure 1). In non-emmetropic eyes the relationship between corneal curvature and axial length was more non-linear than in emmetropes, and axial length was much more strongly related to refractive error, especially in individuals with higher levels of myopia and hypermetropia. Corneal curvature was more strongly associated with refractive error in non-emmetropic eyes than in emmetropic eyes, however the association was markedly weaker than for axial length.

**Figure 1.**
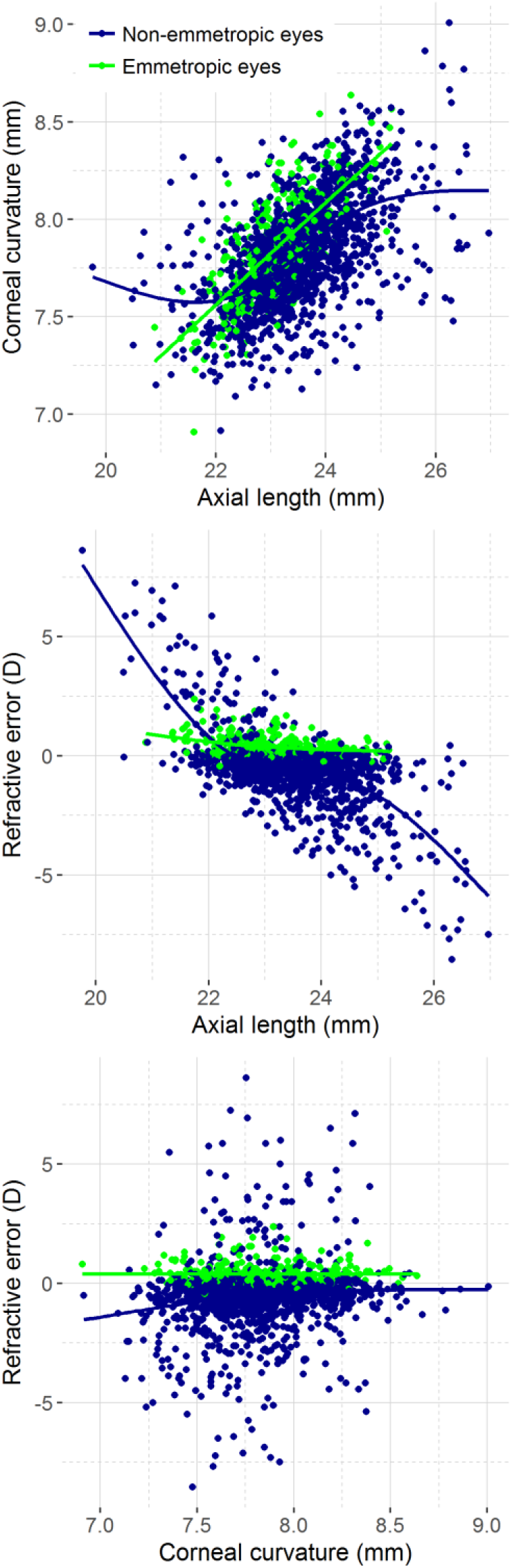
Relationship between corneal curvature and axial length in emmetropic and non-emmetropic eyes of ALSPAC participants. Data are from the emmetropic eye (or eyes) of n=315 individuals with at least 1 emmetropic eye and the eyes of n=1560 individuals in which neither eye was classified as emmetropic. For individuals with both eyes classified as emmetropic, the mean of their 2 eyes was used. (Note that because both sphere and cylinder refractive error were used to classify eyes as emmetropic, some non-emmetropic eyes had a spherical equivalent refractive error that would be within the range typical of emmetropic eyes). All curves were fitted using the default generalized additive model (GAM) function of the ggplot2 geom_smooth function.

### Selection of instrumental variables for eye size in emmetropes

We took advantage of the close (genetically-determined) relationship between corneal curvature and axial length in emmetropes to carry out a GWAS for eye size. Specifically, we carried out a GWAS for corneal curvature in emmetropes in order to identify genetic variants associated with eye size in eyes with optimally scaled ocular components (Figure S3A). This GWAS for corneal curvature in the emmetropic eyes of 22,180 individuals from the UK Biobank cohort led to the identification of 32 independently-associated genetic markers (P <5.0e-08; Table S2). In the independent ALSPAC study sample of 15 year-old children, a polygenic risk score composed of these 32 genetic markers explained approximately 2.5% of the inter-individual variation in both axial length and corneal curvature in emmetropes (Table 1), confirming that this set of markers represents a robust instrumental variable for both corneal curvature and for axial length, i.e. eye size. The 32-marker polygenic risk score was less predictive of eye size – especially for axial length – in children not selected as being emmetropic (Table 1) consistent with the theory that the normal, co-ordinated scaling of ocular component dimensions is disturbed in eyes with myopia or hypermetropia [39, 40].

**Table 1.** Variance in corneal curvature and axial length in ALSPAC participants explained by a polygenic risk score for corneal curvature.

| Sample | Corneal curvature |  |  | Axial length |  |  |
| --- | --- | --- | --- | --- | --- | --- |
|  | N | R<sup>2</sup> | P | N | R<sup>2</sup> | P |
| Emmetropes | 307 | 2.27% | 7.68e-03 | 315 | 2.71% | 2.32e-03 |
| All participants | 1901 | 2.23% | 2.10e-11 | 1909 | 0.66% | 2.11e-04 |

Abbreviations: N=sample size; R<sup>2</sup>=variance explained; P=P-value for polygenic risk score

### Tests for a causal role of eye size in susceptibility to refractive error

Mendelian randomization analysis was carried out using the 32 markers identified in the first stage analysis as instrumental variables, and a combined sample of 139,697 individuals (95,505 from UK Biobank and up to 44,192 from the CREAM consortium) who were *not* selected with regard to being or not being emmetropic as the second stage sample (Figure S3B). This provided strong evidence for a causal role of eye size in determining refractive error (Table 2; Figure 2; Table S3 lists associations between each of the 32 instrumental variables and refractive error in for the UK Biobank sample, the CREAM sample, and the 2 samples combined). A standard inverse-variance weighted (IVW) analysis suggested that genetic predisposition to a 1 mm flatter cornea caused a +1.41 D (95% CI. 0.65 to 2.16) more hypermetropic refractive error (P=2.72e-04). Using the value 2.943 for the ratio of axial length:corneal curvature (see above) this corresponds to a +0.48 D (95% CI. 0.22 to 0.73) more hypermetropic refractive error for an eye with a 1mm longer axial length.

**Figure 2.**
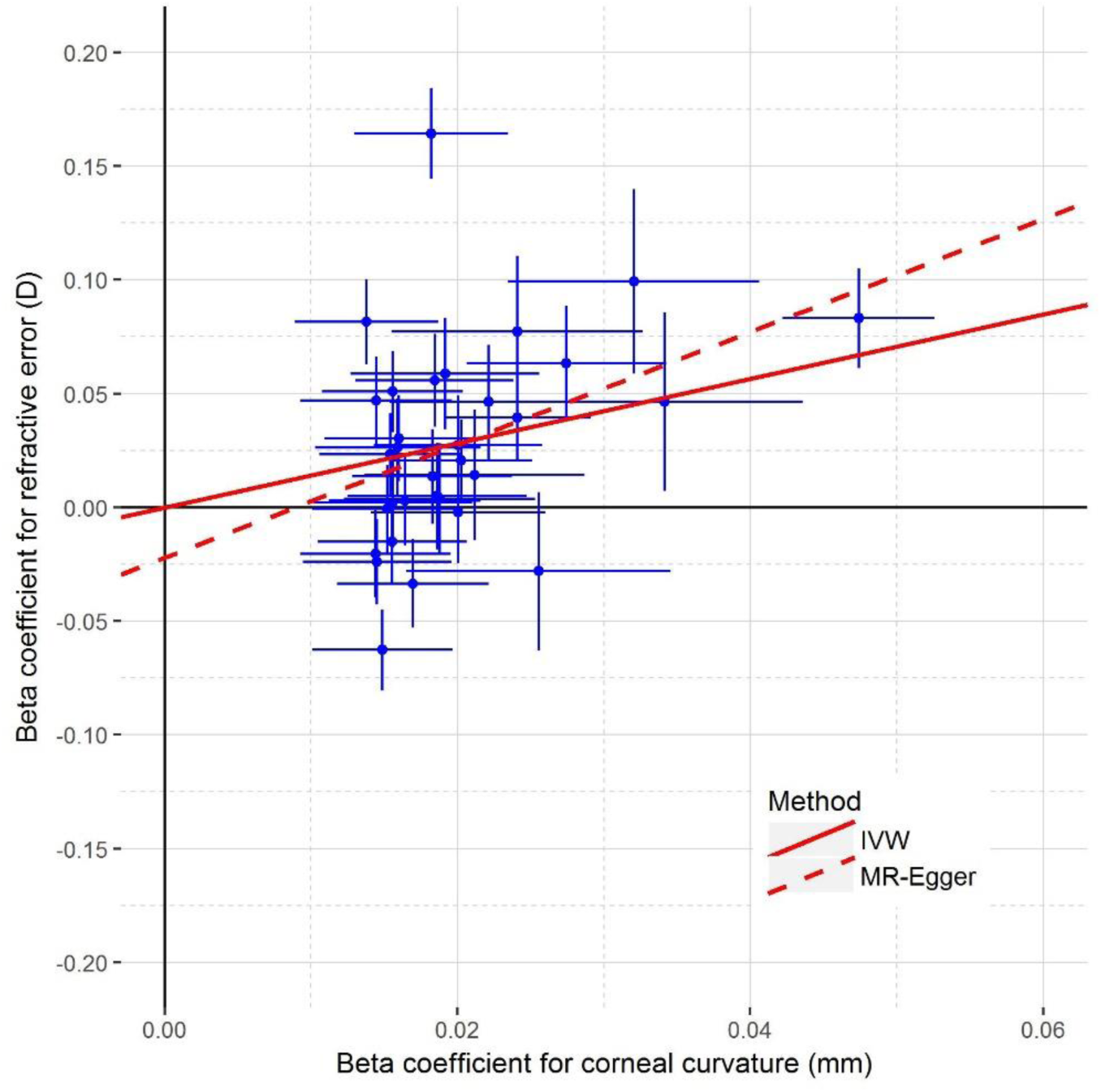
Comparison of estimated effect sizes for association with refractive error and corneal curvature for 32 instrumental variables associated with eye size in emmetropes. Error bars correspond to 95% confidence intervals.

Sensitivity analyses provided additional support for a causal relationship between genetic predisposition for a larger eye size and a more hypermetropic refractive error (Tables 2). Specifically, a simple median-weighted Mendelian randomization causal estimate, which remains valid if up to half of the genetic markers have unwanted pleiotropic effects (i.e. direct effects on refractive error in addition to indirect effects via eye size) and that is resilient against outlier instrumental variables with unusually large or small effects, was +1.36 D for a 1 mm flatter cornea (95% CI. 0.96 to 1.77). An MR-Egger test for directional pleiotropy (here, a tendency for the 32 eye size-associated markers to exhibit direct effects on refractive error consistently in the direction of myopia or consistently in the direction of hypermetropia, irrespective of their influence on eye size) yielded an intercept estimate very close to zero (-0.02 D/mm; 95% CI. -0.07 to 0.03). This suggested that directional pleiotropy was not biasing the causal estimate obtained from convention Mendelian randomisation analysis.

**Table 2.**
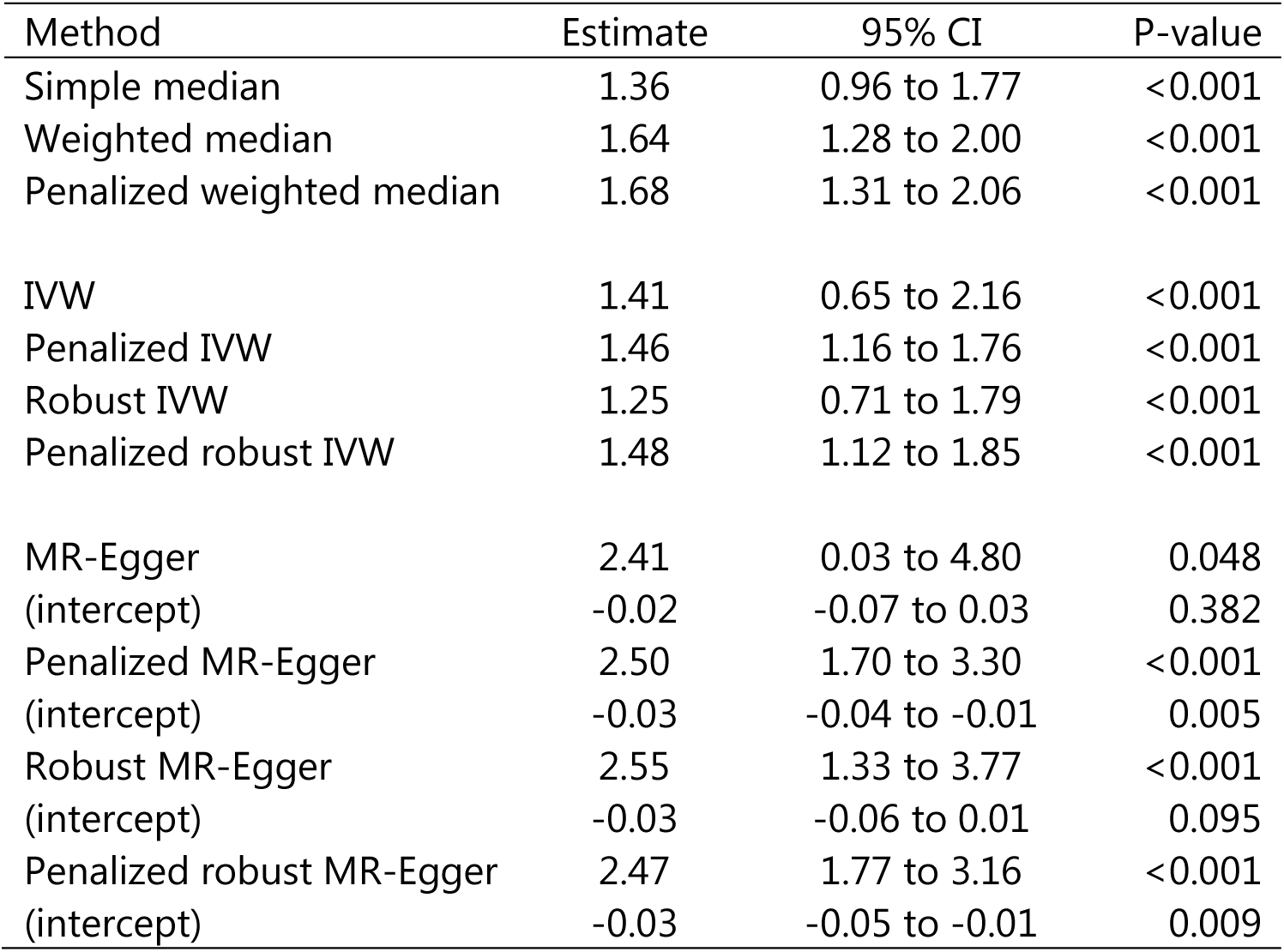
Mendelian randomization analysis for the role of eye size in causing susceptibility to refractive error. Results obtained using the combined UK Biobank and CREAM consortium GWAS analyses as the stage 2 sample. Values are the change in refractive error (D) for a 1mm increase in corneal curvature.

| Method | Estimate | 95% CI | P-value |
| --- | --- | --- | --- |
| Simple median | 1.36 | 0.96 to 1.77 | <0.001 |
| Weighted median | 1.64 | 1.28 to 2.00 | <0.001 |
| Penalized weighted median | 1.68 | 1.31 to 2.06 | <0.001 |
| IVW | 1.41 | 0.65 to 2.16 | <0.001 |
| Penalized IVW | 1.46 | 1.16 to 1.76 | <0.001 |
| Robust IVW | 1.25 | 0.71 to 1.79 | <0.001 |
| Penalized robust IVW | 1.48 | 1.12 to 1.85 | <0.001 |
| MR-Egger | 2.41 | 0.03 to 4.80 | 0.048 |
| (intercept) | -0.02 | -0.07 to 0.03 | 0.382 |
| Penalized MR-Egger | 2.50 | 1.70 to 3.30 | <0.001 |
| (intercept) | -0.03 | -0.04 to -0.01 | 0.005 |
| Robust MR-Egger | 2.55 | 1.33 to 3.77 | <0.001 |
| (intercept) | -0.03 | -0.06 to 0.01 | 0.095 |
| Penalized robust MR-Egger | 2.47 | 1.77 to 3.16 | <0.001 |
| (intercept) | -0.03 | -0.05 to -0.01 | 0.009 |

There was a 16% overlap between our corneal curvature GWAS sample (Mendelian randomization stage 1) and our refractive error GWAS sample (Mendelian randomization stage 2). In the event that instrumental variables are only weakly predictive of the exposure, such sample overlap can bias causal estimates away from zero; so called "weak instrument bias" [41]. Therefore, as a further sensitivity analysis we repeated the Mendelian randomization analyses using only the CREAM consortium refractive error GWAS as the second stage sample. For these analyses, in which there was no overlap between the first and second stage samples, the magnitude and direction of the causal effect estimates were similar to those in the main analyses (Table S4). For example, the IVW causal estimate was + 1.13 D for a 1 mm flatter cornea (95% CI. 0.49 to 1.76) using only the CREAM GWAS results for the second stage (versus +1.41 D/mm when using CREAM plus UK Biobank GWAS results for the second stage).

As with any definition of emmetropia, the definition we adopted (0.00 ≤ SPH ≤ +1.00 D; 0.00 ≤ |CYL| ≤ +1.00 D; VA <0.2 logMAR) was somewhat arbitrary. Therefore, as a further sensitivity analysis, we repeated the corneal curvature GWAS and Mendelian randomisation analysis using an alternative definition [42] of emmetropia: -0.50 ≤ MSE ≤ +0.50 D (along with the requirement for VA <0.2 logMAR); where MSE represents the mean spherical equivalent refractive error. The corneal curvature GWAS using the alternative definition (n=27,569 participants) yielded 38 genetic variants (P<5.0e-08) for use as instrumental variables. The IVW Mendelian randomisation estimate of the causal effect of eye size on refractive error was +1.57 D for a 1 mm flatter cornea (95% CI. 0.96 to 2.18; Table S5), corresponding to +0.53 D more hypermetropia for a 1 mm longer eye (95% CI. 0.33 to 0.74). With the new definition of emmetropia, MR-Egger analysis once again provided no evidence of directional pleiotropy (Egger intercept = -0.01; Table S5). Furthermore, we repeated the GWAS for corneal curvature only in participants (n=12,014) classified as being emmetropic in *both* eyes using the definition -0.50 ≤ MSE ≤ +0.50 D and VA <0.2 logMAR. This identified 12 genetic variants with P<5.0e-08, with a high degree of overlap to those identified above. Mendelian randomisation analysis using these 12 variants as instrumental variables yielded an IVW causal effect estimate of +1.11 D per mm flatter cornea (95% CI. 0.72 to 1.50), which corresponds approximately to a refractive error +0.38 D more hypermetropic per mm longer axial length (95% CI. 0.24 to 0.51). Thus, the causal effect estimate was robust to the exact definition of emmetropia adopted and minimally affected by the 1^st^-stage GWAS in emmetropic eyes being performed in individuals with either at least one eye, or both eyes, classified as emmetropic.

In order to establish whether genetic variants associated with both height (body stature) and eye size were biasing our Mendelian randomisation results – since, for example, height is associated with educational attainment, and this in turn is associated with refractive error [8, 43] – a sensitivity analysis was also carried out using instrumental variables for eye size independent of height (Figure S3A). Thus, the GWAS for corneal curvature was repeated, this time with height included in the analysis model as a continuous covariate. This GWAS yielded 32 genetic variants (P<5.0e-08) for use as instrumental variables (with considerable overlap between the results for GWAS analyses with and without adjustment for height). In the height-adjusted Mendelian randomisation analysis, the IVW estimate of the causal effect of eye size on refractive error was 1.64 D for a 1 mm flatter cornea (95% CI. 0.90 to 2.39; Table S7), corresponding to +0.56 D more hypermetropia for a 1 mm longer eye (95% CI. 0.30 to 0.81). MR-Egger analysis demonstrated no evidence of directional pleiotropy (Egger intercept = -0.01; Table S7). Thus, there was no evidence to suggest that the original causal estimate was biased by pleiotropic effects of the instrumental variables on height.

## Discussion

Previous work has suggested that a larger eye size is a risk factor for myopia. Our Mendelian randomisation findings imply the opposite – namely, that from the perspective of the biological mechanisms acting to optimally scale the human eye, the determinants of normal eye size act such that shorter eyes will tend to be more myopic and larger eyes will tend to be more hypermetropic. Specifically, for each 1mm increase in eye size, our results suggest that the eye is geared towards becoming approximately 0.5 D more hypermetropic.

A key aspect of this study was that genetic variants associated with eye size (i.e. the first stage of Mendelian randomisation) were identified in a sample of individuals selected for emmetropia rather than in the full population. Had such outcome-based selection occurred in the second stage of the Mendelian randomisation, the causal estimate would likely have been affected by collider bias [44]. Crucially, there was no selection of participants based on the outcome variable in the second stage of Mendelian randomisation, thus excluding the possibility of this source of collider bias. Precedents for selection based on the outcome phenotype in the first stage of an analysis include a study by the Emerging Risk Factors Collaboration [45], who identified variants associated with C-Reactive Protein (CRP) in a sample selected for *not* having a history of coronary heart disease (CHD) prior to testing if CRP level is a causal risk factor for CHD, and a study by De Silva et al. [46] who identified variants associated with circulating triglyceride levels in non-diabetics prior to testing if triglyceride levels have a causal role in diabetes.

Our findings have several implications in the context of previous work. Firstly, it seems counterintuitive that a set of genetic variants whose primary role is to generate an eye with correctly scaled ocular components could, at the same time, be "programmed" to link axial and corneal eye growth to hypermetropia. Yet, mild hypermetropia is in fact the norm in most animal populations, in human infants, and in adult humans living in communities not exposed to a modern, westernised environment [47–51], and there is a substantial overlap in the axial length distribution across refractive groups classified as hypermetropes, emmetropes and myopes [38]. Since the visually-guided emmetropisation feedback system is better adapted to up-regulating the rate of axial elongation in eyes that are too hypermetropic (compared to its ability to slow the rate of elongation of eyes that are too myopic) it would be advantageous for the eye to have evolved a tendency towards hypermetropia, not least since there may be a limit to the extent that already-elongated eyes can be remodelled into shorter eyes, whereas the capacity for enlarged eye growth is substantial. Secondly, the result demands an explanation for the negative *phenotypic correlation* between refractive error and axial length that has been reported clinically, instead of the positive correlation predicted by our Mendelian randomisation analysis. Furthermore, this explanation must be able to account for the negative *genetic correlation* between refractive error and axial length that has also been observed [13, 14]. We speculate that the negative phenotypic correlation arises because myopic eyes have axially elongated using distinct molecular pathways to those controlling normal eye growth. This would lead to a breakdown in the usual, carefully balanced scaling of corneal curvature and axial length (and may contribute to the differences in three-dimensional shape between emmetropic and myopic eyes of similar axial length [52, 53]). We further suggest that the observed negative genetic correlation between refractive error and axial length arises because these traits were measured in populations with a high prevalence of myopia; thus, the negative genetic correlation would reflect the effects of genetic variants that lead to an elongated eye that is also a myopic eye. This contrasts with the near zero genetic correlation between refractive error and axial length one might expect in a sample of emmetropic eyes, in which axial length and refractive error would, by definition, be independent. Thus, in a mixed population of emmetropes and myopes, the measured genetic correlation would lie between the zero expected in emmetropes and the high negative value expected in myopes. Thirdly, our results seem to contradict two studies of 6–14 year-old children in which a larger eye size has been shown to be predictive of incident myopia [15, 16]. In one study [15], non-myopic children with myopic parents had longer eyes and less hypermetropic refractions than children without myopic parents, while in the other study [16] children who developed myopia were found to have longer eyes and more myopic refractions 3–4 years before actually being diagnosed as myopic. We suggest that the children with myopic parents [15] and those destined to become myopic [16] were already progressing towards myopia, even though they had not yet reached the -0.75 D threshold level used by the two studies' authors to define myopic status. Therefore the normal scaling of the ocular components of these children – and the causal link between longer eyes and a more hypermetropic refractive error suggested by our Mendelian randomisation analysis – would have been offset by the genetic and environmental risk factors causing the breakdown of this balanced scaling as the children developed myopia. Finally, our findings raise the idea of novel approach for slowing the progression of myopia, based on exploiting the causal link between a larger eye size and greater hypermetropia. If a drug capable of up-regulating a genetic pathway controlling eye size was available, then it should – at least in theory – both increase eye size and make the eye less myopic. However, despite any appeal of such an approach, we caution that it would also pose risks. The likelihood of pathological complications in myopic eyes correlates with axial length [1] and therefore even if an eye size-based treatment intervention successfully flattened the curvature of the cornea and reduced the degree of myopia, the treatment's effect of increasing axial eye length could nevertheless put the eye at greater risk of pathology.

This study identified 32 genetic variants associated with eye size, of which 30 implicate novel loci (the *RSPO1* and *PDGFRA* loci have been associated with larger eye size in previous work [54, 55]). The list of the nearest genes at the top loci (Table S2) includes genes associated with spherical refractive error *(PRSS56* [11]), astigmatism *(PDGFRA, LINC00340* [56]) and exfoliation glaucoma *(LOXL1* [57]), as well as 2 members of the *ADAMTS* family.

Strengths of this work were that it took advantage of the only large sample of emmetropes with genotype information currently available worldwide (n=22,180) and leveraged information on refractive error from the largest datasets available (total n=139,697), thus providing precise effect size estimates. Furthermore, while previous observational studies have reported conflicting descriptions of the relationship between eye size and refractive error, likely due to the diverse age ranges and myopia prevalence rates of their study cohorts, here we sought to provide a definitive assessment of the *causal* relationship between eye size in emmetropes and refractive error, operating across the life course. The major limitations of the work are the two central assumptions inherent in Mendelian randomisation studies: (1) that the instrumental variables (eye size SNPs) only exert effects on the outcome (refractive error) via the exposure (eye size) and not directly, and (2) the instrumental variables do not exert effects on confounders of the exposure-outcome relationship. The MR-Egger sensitivity analysis designed to test for directional pleiotropy, i.e. invalidation of the first assumption in such a way as to bias our causal estimate, suggested that directional pleiotropy was essentially absent. A prior study [58] has provided evidence that the second assumption is generally valid, by showing that – apart from rare exceptions – the transmission of alleles of instrumental variable SNPs is independent of the levels of common confounders such as age, socioeconomic status, and body weight.

## Conclusion

Past studies have provided conflicting views regarding whether eye size early in life is a risk factor for myopia [15–17], and whether genetic variants contributing to normal variation in eye size predispose individuals to myopia [13, 14, 18, 54]. Here, for the first time, we explicitly test the hypothesis that a larger eye size is a *causal risk factor* for myopia. Our results provide strong evidence against the hypothesis, and instead suggest that each 1 mm increase in eye length is associated with a +0.48 D (95% CI. 0.22 to 0.73; P<0.001) more hypermetropic (and thus *less* myopic) refractive error. We argue that the conflicting evidence for a relationship between larger eye size and incident myopia can be explained by past choices of study sample: in studies with a high proportion of participants destined to become myopic, an observational association between eye size and myopia will arise because an abnormal degree of axial elongation will have already occured in eyes developing myopia even before they meet the criteria for classifying an eye as myopic. Crucially, our findings imply that the molecular pathways controlling normal variation in eye size are distinct from those used to increase the axial length of the eye during myopia development.

## Acknowledgements

<u>UK Biobank:</u> This research has been conducted using the UK Biobank Resource (applications #17351 and #17615).

<u>ALSPAC:</u> We are extremely grateful to all the ALSPAC families who took part in this study, the midwives for their help in recruiting them, and the whole ALSPAC team, which includes interviewers, computer and laboratory technicians, clerical workers, research scientists, volunteers, managers, receptionists and nurses.

<u>Infrastructure:</u> Data analysis was carried out using the RAVEN computing cluster, maintained by the ARCCA group at Cardiff University ARCCA and the BLUE CRYSTAL3 computing cluster maintained by the HPC group at the University of Bristol.

<u>Funding:</u> This research was specifically funded by NIHR Senior Research Fellowship award SRF-2015-08-005, the Global Education Program of the Russian Federation government, and the National Eye Research Centre grant SAC015. The UK Medical Research Council and the Wellcome Trust (Grant ref: 102215/2/13/2) and the University of Bristol provide core support for ALSPAC. ALSPAC GWAS data was generated by Sample Logistics and Genotyping Facilities at the Wellcome Trust Sanger Institute and LabCorp (Laboratory Corporation of America) using support from 23andMe.

## Supplementary Information

**Table S1.**
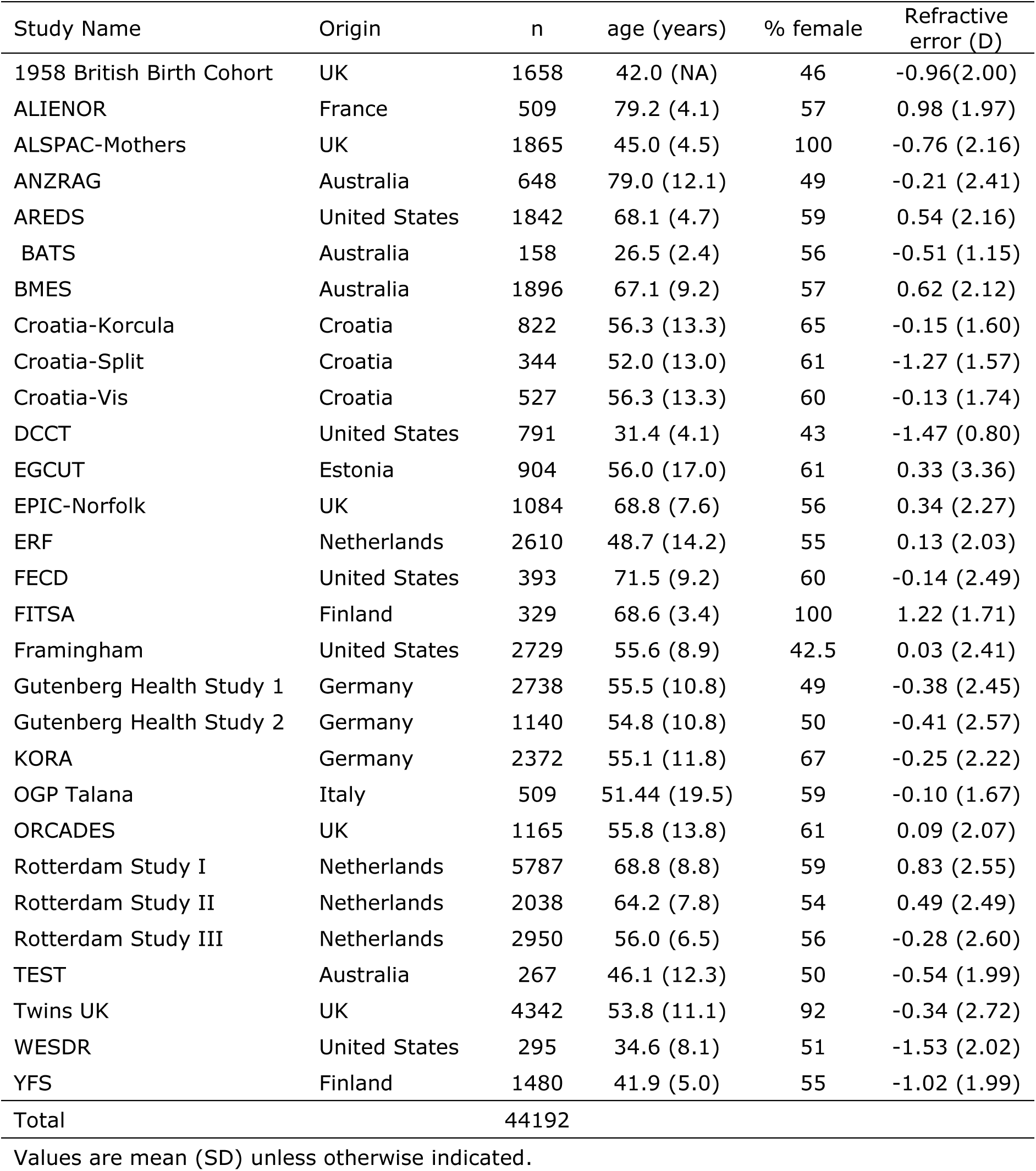
Demographic characteristics of the CREAM consortium study cohorts.

| Study Name | Origin | n | age (years) | % female | Refractive error (D) |
| --- | --- | --- | --- | --- | --- |
| 1958 British Birth Cohort | UK | 1658 | 42.0 (NA) | 46 | -0.96(2.00) |
| ALIENOR | France | 509 | 79.2 (4.1) | 57 | 0.98 (1.97) |
| ALSPAC-Mothers | UK | 1865 | 45.0 (4.5) | 100 | -0.76 (2.16) |
| ANZRAG | Australia | 648 | 79.0 (12.1) | 49 | -0.21 (2.41) |
| AREDS | United States | 1842 | 68.1 (4.7) | 59 | 0.54 (2.16) |
| BATS | Australia | 158 | 26.5 (2.4) | 56 | -0.51 (1.15) |
| BMES | Australia | 1896 | 67.1 (9.2) | 57 | 0.62 (2.12) |
| Croatia-Korcula | Croatia | 822 | 56.3 (13.3) | 65 | -0.15 (1.60) |
| Croatia-Split | Croatia | 344 | 52.0 (13.0) | 61 | -1.27 (1.57) |
| Croatia-Vis | Croatia | 527 | 56.3 (13.3) | 60 | -0.13 (1.74) |
| DCCT | United States | 791 | 31.4 (4.1) | 43 | -1.47 (0.80) |
| EGCUT | Estonia | 904 | 56.0 (17.0) | 61 | 0.33 (3.36) |
| EPIC-Norfolk | UK | 1084 | 68.8 (7.6) | 56 | 0.34 (2.27) |
| ERF | Netherlands | 2610 | 48.7 (14.2) | 55 | 0.13 (2.03) |
| FECD | United States | 393 | 71.5 (9.2) | 60 | -0.14 (2.49) |
| FITSA | Finland | 329 | 68.6 (3.4) | 100 | 1.22 (1.71) |
| Framingham | United States | 2729 | 55.6 (8.9) | 42.5 | 0.03 (2.41) |
| Gutenberg Health Study 1 | Germany | 2738 | 55.5 (10.8) | 49 | -0.38 (2.45) |
| Gutenberg Health Study 2 | Germany | 1140 | 54.8 (10.8) | 50 | -0.41 (2.57) |
| KORA | Germany | 2372 | 55.1 (11.8) | 67 | -0.25 (2.22) |
| OGP Talana | Italy | 509 | 51.44 (19.5) | 59 | -0.10 (1.67) |
| ORCADES | UK | 1165 | 55.8 (13.8) | 61 | 0.09 (2.07) |
| Rotterdam Study I | Netherlands | 5787 | 68.8 (8.8) | 59 | 0.83 (2.55) |
| Rotterdam Study II | Netherlands | 2038 | 64.2 (7.8) | 54 | 0.49 (2.49) |
| Rotterdam Study III | Netherlands | 2950 | 56.0 (6.5) | 56 | -0.28 (2.60) |
| TEST | Australia | 267 | 46.1 (12.3) | 50 | -0.54 (1.99) |
| Twins UK | UK | 4342 | 53.8 (11.1) | 92 | -0.34 (2.72) |
| WESDR | United States | 295 | 34.6 (8.1) | 51 | -1.53 (2.02) |
| YFS | Finland | 1480 | 41.9 (5.0) | 55 | -1.02 (1.99) |
| Total |  | 44192 |  |  |  |

Values are mean (SD) unless otherwise indicated.

**Table S2.**
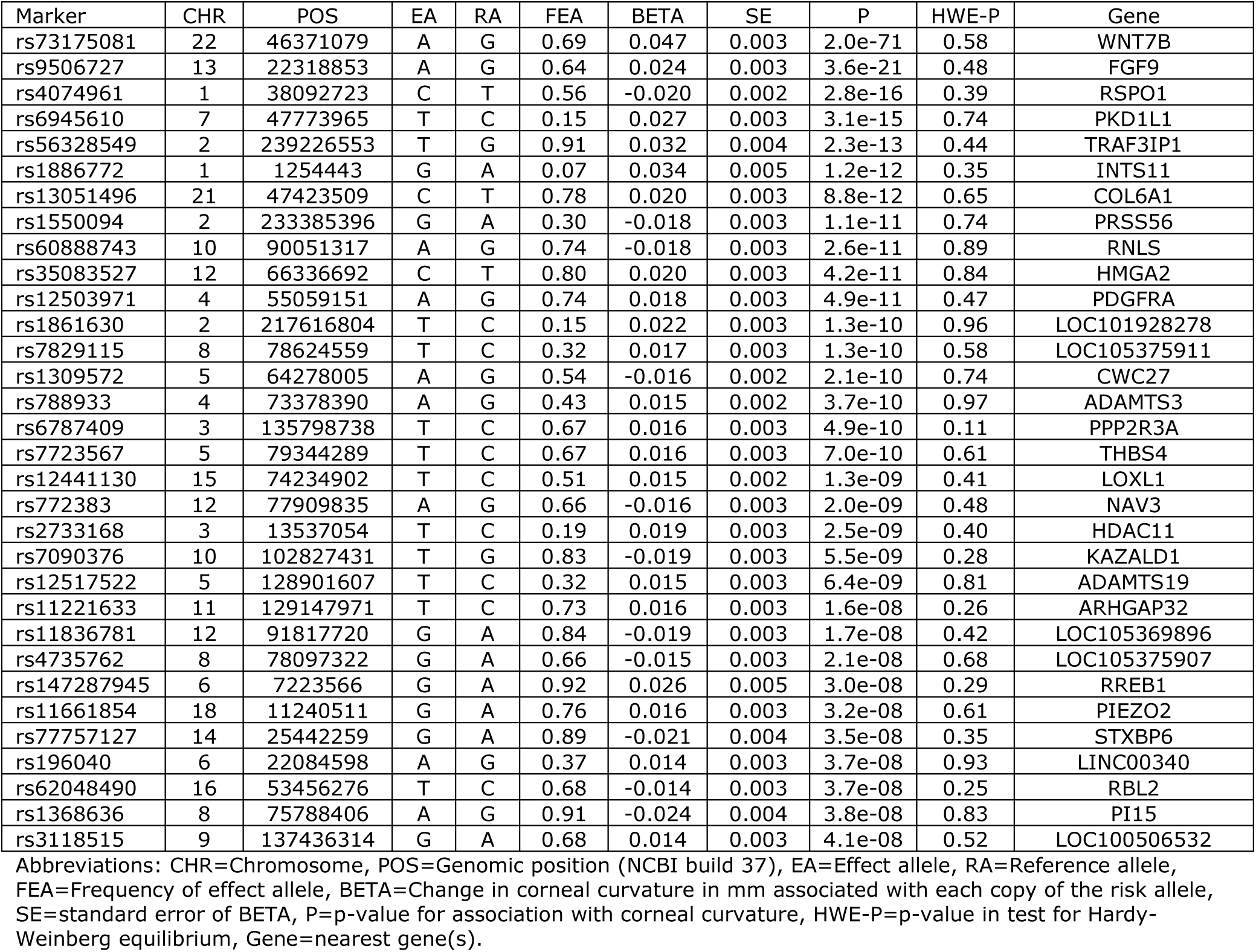
Instrumental variables for eye size in emmetropes: Genetic markers associated with corneal curvature in emmetropes from UK Biobank (n=22,180).

**Table S3.** Stage 2 Mendelian randomization results. The association of the 32 instrumental variables with refractive error in the UK Biobank GWAS, the CREAM consortium GWAS meta-analysis, and the combined sample.

| SNP | CHR | POS | EA | RA | CREAM |  |  |  | UK Biobank |  |  |  | Combined sample |  |  |  |
| --- | --- | --- | --- | --- | --- | --- | --- | --- | --- | --- | --- | --- | --- | --- | --- | --- |
|  |  |  |  |  | BETA | SE | P | N | BETA | SE | P | N | BETA | SE | P | N |
| rs1886772 | 1 | 1254443 | G | A | 0.015 | 0.041 | 7.09E-01 | 24639 | 0.056 | 0.023 | 3.30E-02 | 95505 | 0.046 | 0.02 | 2.04E-02 | 120144 |
| rs4074961 | 1 | 38092723 | T | C | 0.008 | 0.015 | 6.16E-01 | 43925 | 0.029 | 0.012 | 1.30E-02 | 95505 | 0.021 | 0.009 | 2.67E-02 | 139430 |
| rs1861630 | 2 | 217616804 | T | C | 0.04 | 0.021 | 5.19E-02 | 43924 | 0.05 | 0.016 | 3.00E-03 | 95505 | 0.046 | 0.013 | 2.94E-04 | 139429 |
| rs1550094 | 2 | 233385396 | A | G | 0.108 | 0.017 | 1.31E-10 | 43197 | 0.196 | 0.013 | 4.10E-59 | 95505 | 0.164 | 0.01 | 5.08E-60 | 138702 |
| rs56328549 | 2 | 239226553 | T | G | 0.022 | 0.027 | 4.02E-01 | 43912 | 0.098 | 0.021 | 8.30E-06 | 95505 | 0.069 | 0.016 | 2.31E-05 | 139417 |
| rs2733168 | 3 | 13537054 | T | C | 0.004 | 0.02 | 8.43E-01 | 43925 | 0.006 | 0.015 | 9.60E-01 | 95505 | 0.005 | 0.012 | 6.77E-01 | 139430 |
| rs6787409 | 3 | 135798738 | T | C | -0.01 | 0.016 | 5.31E-01 | 43886 | 0.011 | 0.012 | 3.30E-01 | 95505 | 0.003 | 0.01 | 7.77E-01 | 139391 |
| rs12503971 | 4 | 55059151 | A | G | 0.018 | 0.018 | 3.07E-01 | 43229 | 0.011 | 0.013 | 4.80E-01 | 95505 | 0.014 | 0.011 | 1.98E-01 | 138734 |
| rs788933 | 4 | 73378390 | A | G | 0.039 | 0.015 | 1.00E-02 | 43925 | 0.014 | 0.012 | 1.60E-01 | 95505 | 0.023 | 0.009 | 1.16E-02 | 139430 |
| rs1309572 | 5 | 64278005 | G | A | 0.059 | 0.015 | 6.34E-05 | 43925 | 0.046 | 0.012 | 5.80E-05 | 95505 | 0.051 | 0.009 | 2.51E-08 | 139430 |
| rs7723567 | 5 | 79344289 | T | C | 0.015 | 0.016 | 3.37E-01 | 43920 | 0.04 | 0.012 | 2.00E-03 | 95505 | 0.03 | 0.01 | 1.67E-03 | 139425 |
| rs12517522 | 5 | 128901607 | T | C | 0.003 | 0.016 | 8.78E-01 | 43911 | -0.003 | 0.012 | 8.80E-01 | 95505 | -0.001 | 0.01 | 9.41E-01 | 139416 |
| rs147287945 | 6 | 7223566 | G | A | -0.019 | 0.031 | 5.38E-01 | 39926 | -0.032 | 0.022 | 2.30E-01 | 95505 | -0.028 | 0.018 | 1.14E-01 | 135431 |
| rs196040 | 6 | 22084598 | A | G | 0.08 | 0.015 | 1.83E-07 | 43904 | 0.082 | 0.012 | 9.40E-13 | 95505 | 0.082 | 0.01 | 8.74E-18 | 139409 |
| rs6945610 | 7 | 47773965 | T | C | 0.046 | 0.021 | 2.35E-02 | 43918 | 0.074 | 0.017 | 4.10E-05 | 95505 | 0.063 | 0.013 | 9.20E-07 | 139423 |
| rs1368636 | 8 | 75788406 | G | A | 0.08 | 0.029 | 6.28E-03 | 43839 | 0.076 | 0.021 | 5.50E-05 | 95505 | 0.077 | 0.017 | 5.29E-06 | 139344 |
| rs4735762 | 8 | 78097322 | A | G | -0.014 | 0.015 | 3.69E-01 | 43923 | -0.031 | 0.012 | 1.60E-02 | 95505 | -0.024 | 0.01 | 1.22E-02 | 139428 |
| rs7829115 | 8 | 78624559 | T | C | -0.023 | 0.016 | 1.47E-01 | 43858 | -0.04 | 0.013 | 1.10E-03 | 95505 | -0.034 | 0.01 | 6.97E-04 | 139363 |
| rs3118515 | 9 | 137436314 | G | A | 0.041 | 0.016 | 1.09E-02 | 43925 | 0.051 | 0.012 | 2.50E-05 | 95505 | 0.047 | 0.01 | 1.72E-06 | 139430 |
| rs60888743 | 10 | 90051317 | G | A | 0.046 | 0.017 | 6.90E-03 | 43924 | 0.062 | 0.013 | 5.40E-07 | 95505 | 0.056 | 0.01 | 8.70E-08 | 139429 |
| rs7090376 | 10 | 102827431 | G | T | 0.041 | 0.021 | 5.04E-02 | 43925 | 0.069 | 0.016 | 8.30E-06 | 95505 | 0.059 | 0.013 | 2.57E-06 | 139430 |
| rs11221633 | 11 | 129147971 | T | C | 0.002 | 0.017 | 9.25E-01 | 43916 | 0.002 | 0.013 | 5.80E-01 | 95505 | 0.002 | 0.01 | 8.38E-01 | 139421 |
| rs35083527 | 12 | 66336692 | C | T | -0.021 | 0.018 | 2.52E-01 | 43924 | 0.009 | 0.014 | 8.10E-01 | 95505 | -0.002 | 0.011 | 8.44E-01 | 139429 |
| rs772383 | 12 | 77909835 | G | A | 0.002 | 0.015 | 9.06E-01 | 43925 | -0.026 | 0.012 | 3.10E-02 | 95505 | -0.015 | 0.01 | 1.12E-01 | 139430 |
| rs11836781 | 12 | 91817720 | A | G | 0.006 | 0.02 | 7.78E-01 | 43925 | 0.002 | 0.016 | 7.50E-01 | 95505 | 0.004 | 0.013 | 7.70E-01 | 139430 |
| rs9506727 | 13 | 22318853 | A | G | 0.033 | 0.016 | 3.52E-02 | 43885 | 0.044 | 0.012 | 1.50E-04 | 95505 | 0.04 | 0.01 | 3.36E-05 | 139390 |
| rs77757127 | 14 | 25442259 | A | G | -0.013 | 0.024 | 5.99E-01 | 40045 | 0.029 | 0.018 | 7.50E-02 | 95505 | 0.014 | 0.015 | 3.33E-01 | 135550 |
| rs12441130 | 15 | 74234902 | T | C | -0.053 | 0.015 | 3.39E-04 | 43925 | -0.069 | 0.012 | 1.80E-09 | 95505 | -0.063 | 0.009 | 5.91E-12 | 139430 |
| rs62048490 | 16 | 53456276 | C | T | 0.013 | 0.016 | 4.07E-01 | 43917 | -0.042 | 0.012 | 2.90E-04 | 95505 | -0.021 | 0.01 | 3.49E-02 | 139422 |
| rs11661854 | 18 | 11240511 | G | A | 0.039 | 0.018 | 2.75E-02 | 43913 | 0.018 | 0.014 | 1.20E-01 | 95505 | 0.026 | 0.011 | 1.58E-02 | 139418 |
| rs13051496 | 21 | 47423509 | C | T | 0.019 | 0.019 | 3.20E-01 | 43904 | 0.032 | 0.014 | 2.10E-02 | 95505 | 0.027 | 0.011 | 1.39E-02 | 139409 |
| rs73175081 | 22 | 46371079 | A | G | 0.07 | 0.025 | 4.75E-03 | 24448 | 0.086 | 0.013 | 2.60E-11 | 95505 | 0.083 | 0.011 | 1.18E-13 | 119953 |

**Table S4.** Mendelian randomization analysis for the role of eye size in causing susceptibility to refractive error, using non-overlapping samples in the first stage (UK Biobank emmetropes) and second stage (CREAM consortium cohorts). Values are estimates of the causal effect on refractive error (D) of a 1mm increase in corneal curvature.

| Method | Estimate | 95% CI | P-value |
| --- | --- | --- | --- |
| Simple median | 0.91 | 0.39 to 1.44 | 0.001 |
| Weighted median | 0.97 | 0.45 to 1.49 | <0.001 |
| Penalized weighted median | 0.96 | 0.43 to 1.48 | <0.001 |
| IVW | 1.13 | 0.49 to 1.76 | 0.001 |
| Penalized IVW | 0.92 | 0.51 to 1.32 | <0.001 |
| Robust IVW | 1.04 | 0.51 to 1.57 | 0.000 |
| Penalized robust IVW | 0.93 | 0.50 to 1.36 | <0.001 |
| MR-Egger | 1.26 | -1.05 to 3.57 | 0.285 |
| (intercept) | 0.00 | -0.05 to 0.04 | 0.906 |
| Penalized MR-Egger | 1.51 | 0.05 to 2.97 | 0.043 |
| (intercept) | -0.01 | -0.04 to 0.02 | 0.426 |
| Robust MR-Egger | 1.37 | -0.08 to 2.81 | 0.064 |
| (intercept) | -0.01 | -0.05 to 0.03 | 0.721 |
| Penalized robust MR-Egger | 1.57 | 0.59 to 2.55 | 0.002 |
| (intercept) | -0.01 | -0.04 to 0.01 | 0.296 |

**Table S5.**
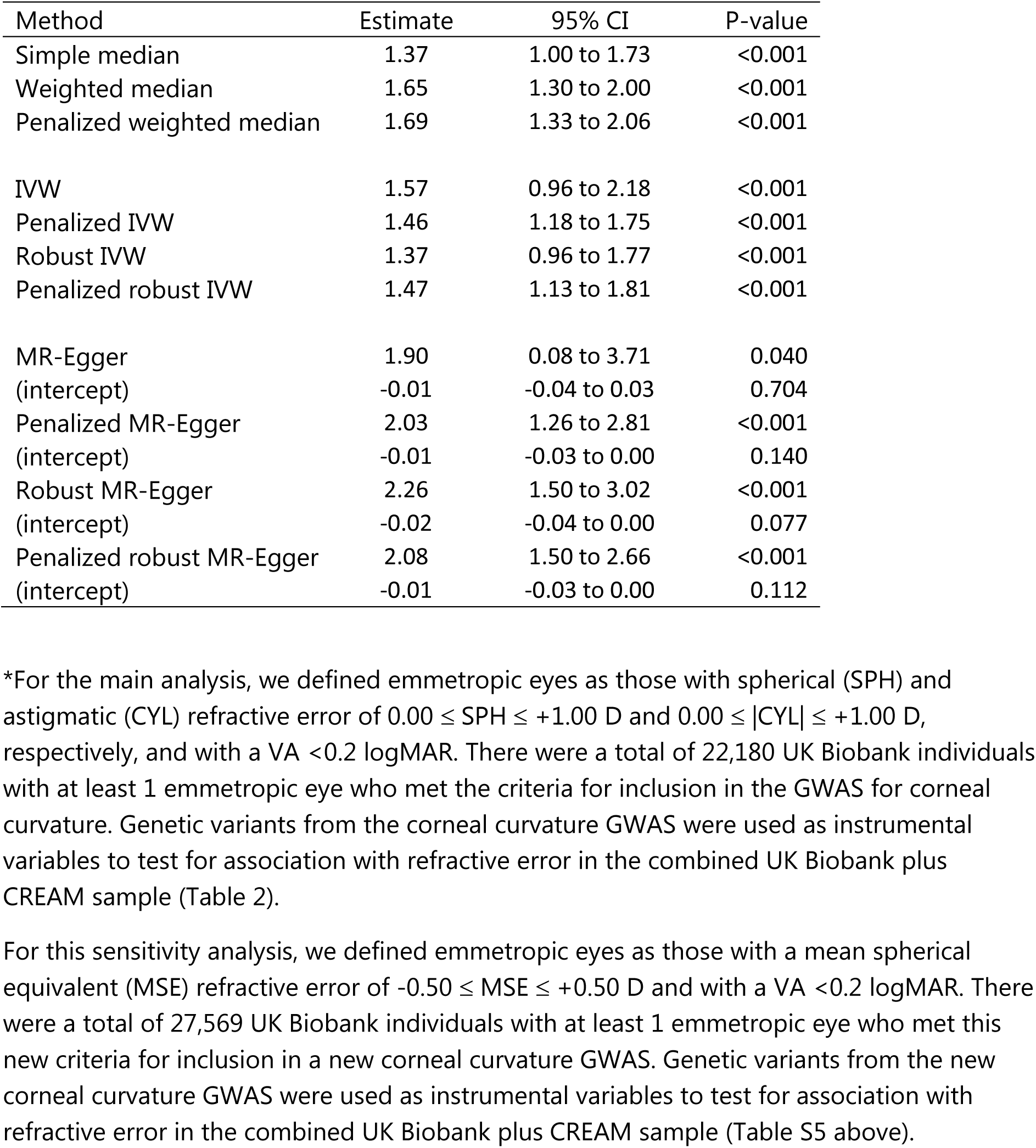
Mendelian randomization analysis for the role of eye size in causing susceptibility to refractive error, using an alternative definition of "emmetropia". Values are estimates of the causal effect on refractive error (D) of a 1mm increase in corneal curvature.

| Method | Estimate | 95% CI | P-value |
| --- | --- | --- | --- |
| Simple median | 1.37 | 1.00 to 1.73 | <0.001 |
| Weighted median | 1.65 | 1.30 to 2.00 | <0.001 |
| Penalized weighted median | 1.69 | 1.33 to 2.06 | <0.001 |
| IVW | 1.57 | 0.96 to 2.18 | <0.001 |
| Penalized IVW | 1.46 | 1.18 to 1.75 | <0.001 |
| Robust IVW | 1.37 | 0.96 to 1.77 | <0.001 |
| Penalized robust IVW | 1.47 | 1.13 to 1.81 | <0.001 |
| MR-Egger | 1.90 | 0.08 to 3.71 | 0.040 |
| (intercept) | -0.01 | -0.04 to 0.03 | 0.704 |
| Penalized MR-Egger | 2.03 | 1.26 to 2.81 | <0.001 |
| (intercept) | -0.01 | -0.03 to 0.00 | 0.140 |
| Robust MR-Egger | 2.26 | 1.50 to 3.02 | <0.001 |
| (intercept) | -0.02 | -0.04 to 0.00 | 0.077 |
| Penalized robust MR-Egger | 2.08 | 1.50 to 2.66 | <0.001 |
| (intercept) | -0.01 | -0.03 to 0.00 | 0.112 |

\*For the main analysis, we defined emmetropic eyes as those with spherical (SPH) and astigmatic (CYL) refractive error of $0.00 \leq \text{SPH} \leq +1.00$ D and $0.00 \leq |\text{CYL}| \leq +1.00$ D, respectively, and with a VA <0.2 logMAR. There were a total of 22,180 UK Biobank individuals with at least 1 emmetropic eye who met the criteria for inclusion in the GWAS for corneal curvature. Genetic variants from the corneal curvature GWAS were used as instrumental variables to test for association with refractive error in the combined UK Biobank plus CREAM sample (Table 2).
For this sensitivity analysis, we defined emmetropic eyes as those with a mean spherical equivalent (MSE) refractive error of $-0.50 \leq \text{MSE} \leq +0.50$ D and with a VA <0.2 logMAR. There were a total of 27,569 UK Biobank individuals with at least 1 emmetropic eye who met this new criteria for inclusion in a new corneal curvature GWAS. Genetic variants from the new corneal curvature GWAS were used as instrumental variables to test for association with refractive error in the combined UK Biobank plus CREAM sample (Table S5 above).

**Table S6.** Mendelian randomization analysis for the role of eye size in causing susceptibility to refractive error, using as the 1^st^ stage a GWAS for corneal curvature in participants classified as emmetropic in both eyes. Emmetropia was defined in for Table S5. Values are estimates of the causal effect on refractive error (D) of a 1mm increase in corneal curvature.

| Method | Estimate | 95% CI | P-value |
| --- | --- | --- | --- |
| Simple median | 0.91 | 0.48 to 1.34 | <0.001 |
| Weighted median | 1.19 | 0.76 to 1.62 | <0.001 |
| Penalized weighted median | 0.87 | 0.46 to 1.28 | <0.001 |
| IVW | 1.11 | 0.72 to 1.50 | <0.001 |
| Penalized IVW | 0.90 | 0.54 to 1.25 | <0.001 |
| Robust IVW | 1.08 | 0.54 to 1.61 | <0.001 |
| Penalized robust IVW | 0.88 | 0.49 to 1.28 | <0.001 |
| MR-Egger | 1.73 | 0.45 to 3.00 | 0.008 |
| (intercept) | -0.02 | -0.05 to 0.02 | 0.317 |
| Penalized MR-Egger | 1.73 | 0.45 to 3.00 | 0.008 |
| (intercept) | -0.02 | -0.05 to 0.02 | 0.317 |
| Robust MR-Egger | 1.73 | 0.01 to 3.45 | 0.049 |
| (intercept) | -0.02 | -0.06 to 0.02 | 0.407 |
| Penalized robust MR-Egger | 1.73 | 0.01 to 3.45 | 0.049 |
| (intercept) | -0.02 | -0.06 to 0.02 | 0.407 |

**Table S7.** Mendelian randomization analysis for the role of eye size in causing susceptibility to refractive error, using as the 1^st^ stage a GWAS for corneal curvature with height as a covariate (i.e. eye size independent of body size). Values are estimates of the causal effect on refractive error (D) of a 1mm increase in corneal curvature.

| Method | Estimate | 95% CI | P-value |
| --- | --- | --- | --- |
| Simple median | 1.48 | 1.09 to 1.86 | <0.001 |
| Weighted median | 1.68 | 1.31 to 2.05 | <0.001 |
| Penalized weighted median | 1.71 | 1.33 to 2.09 | <0.001 |
| IVW | 1.64 | 0.90 to 2.39 | <0.001 |
| Penalized IVW | 1.60 | 1.28 to 1.92 | <0.001 |
| Robust IVW | 1.52 | 1.06 to 1.98 | <0.001 |
| Penalized robust IVW | 1.61 | 1.28 to 1.94 | <0.001 |
| MR-Egger | 2.10 | -0.28 to 4.47 | 0.083 |
| (intercept) | -0.01 | -0.06 to 0.04 | 0.692 |
| Penalized MR-Egger | 2.09 | 1.12 to 3.06 | <0.001 |
| (intercept) | -0.01 | -0.03 to 0.01 | 0.330 |
| Robust MR-Egger | 2.10 | 1.12 to 3.09 | <0.001 |
| (intercept) | -0.01 | -0.04 to 0.02 | 0.371 |
| Penalized robust MR-Egger | 2.10 | 1.49 to 2.71 | <0.001 |
| (intercept) | -0.01 | -0.03 to 0.01 | 0.269 |

**Table S8.**
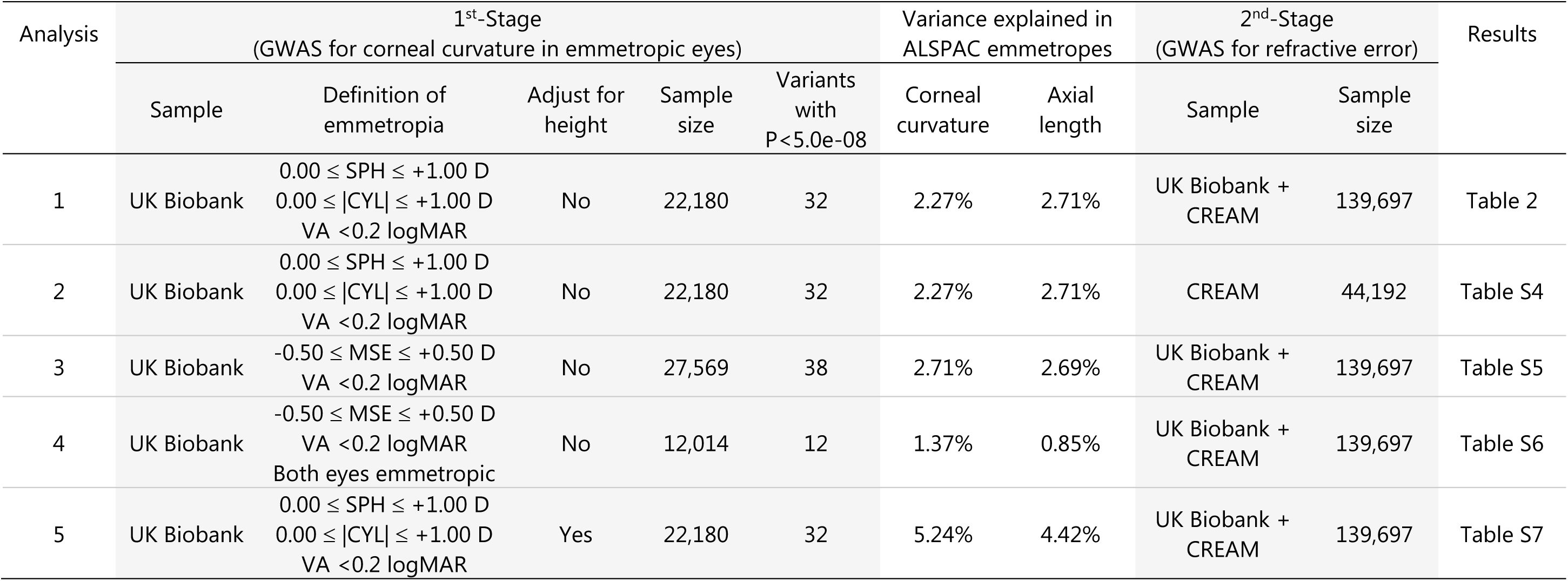
Summary of analyses.

**Figure S1.**
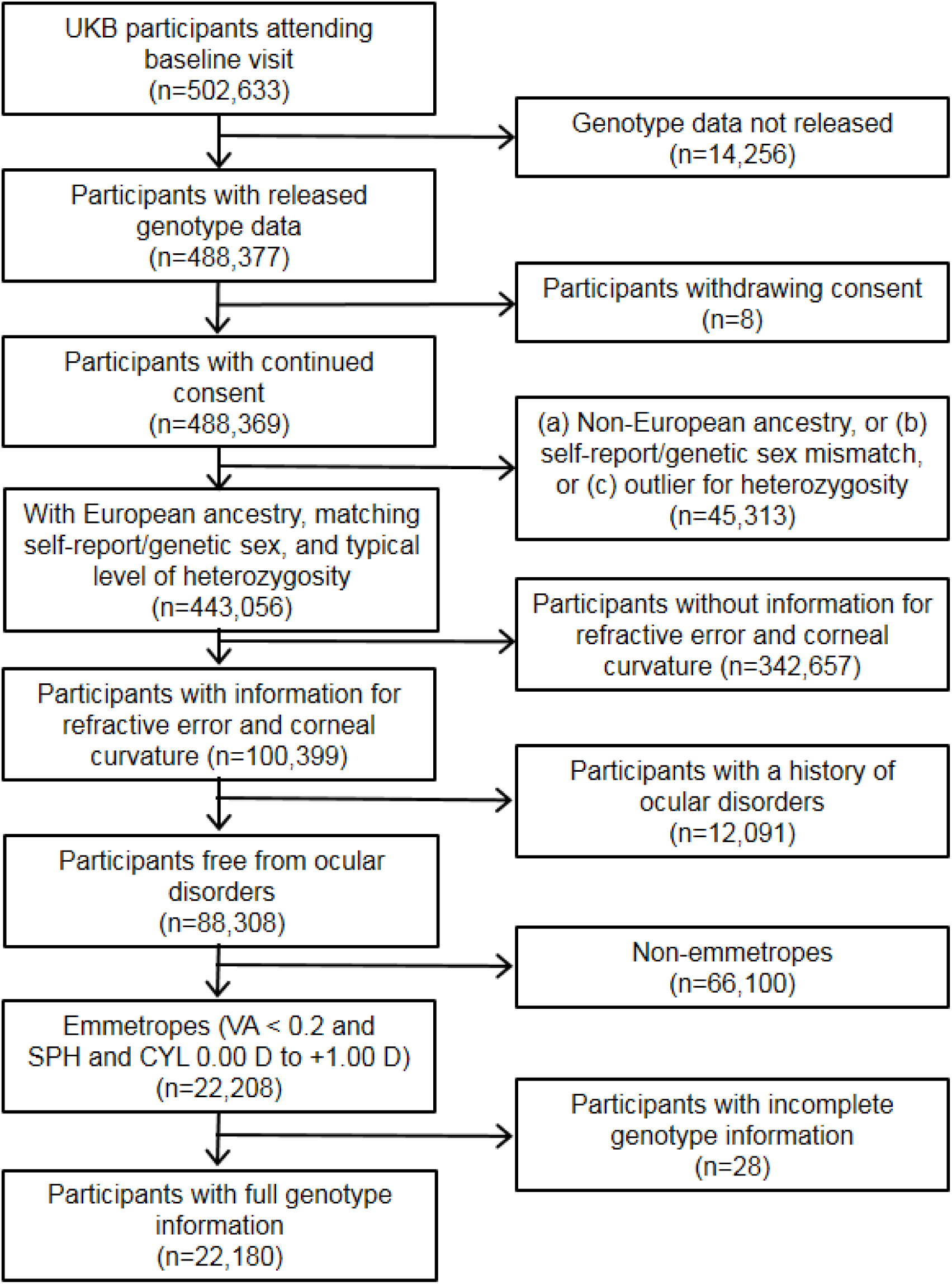
Selection of UK Biobank emmetropic participants for corneal curvature GWAS.

**Figure S2.**
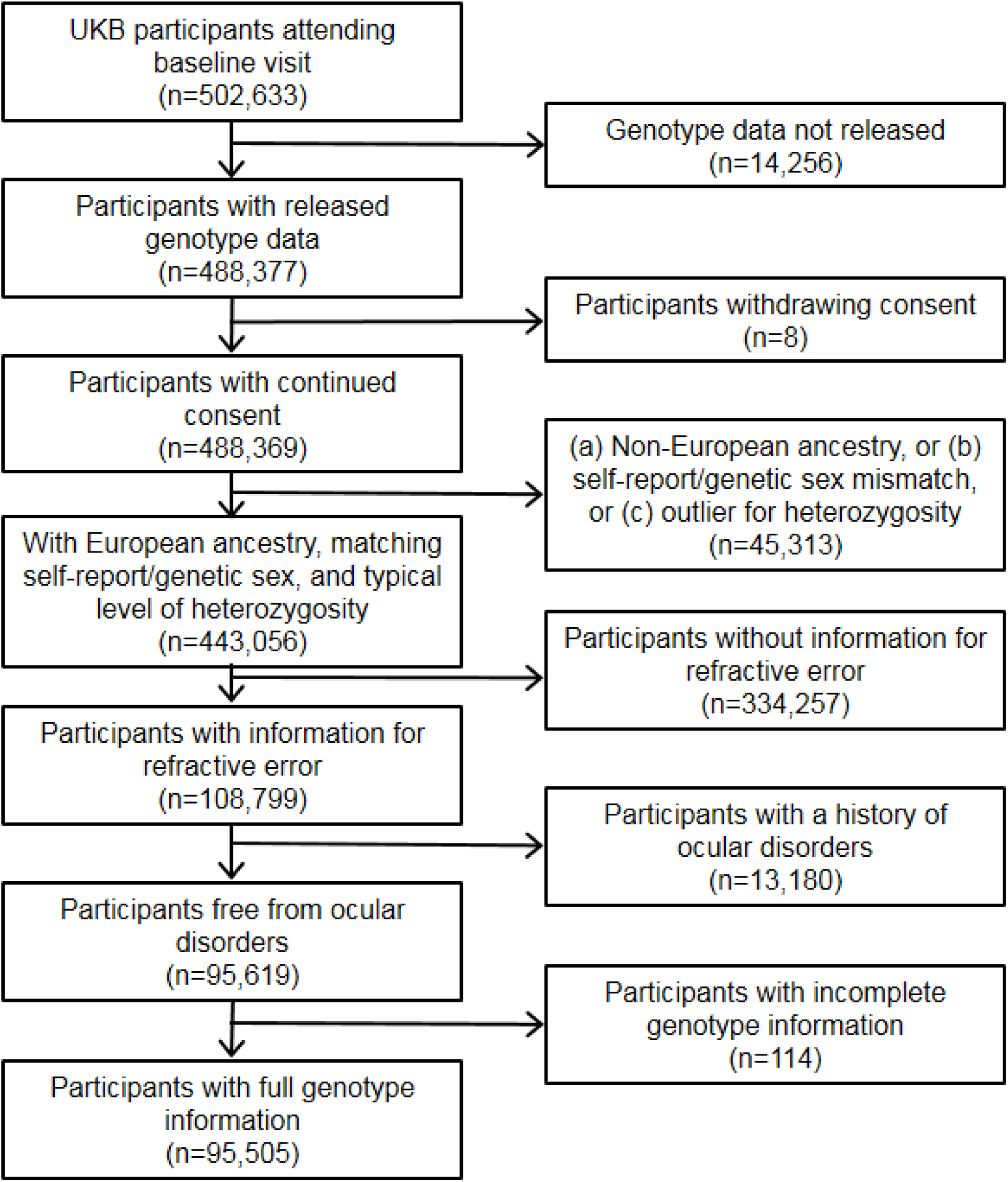
Selection of UK Biobank participants for the refractive error GWAS.

**Figure S3.**
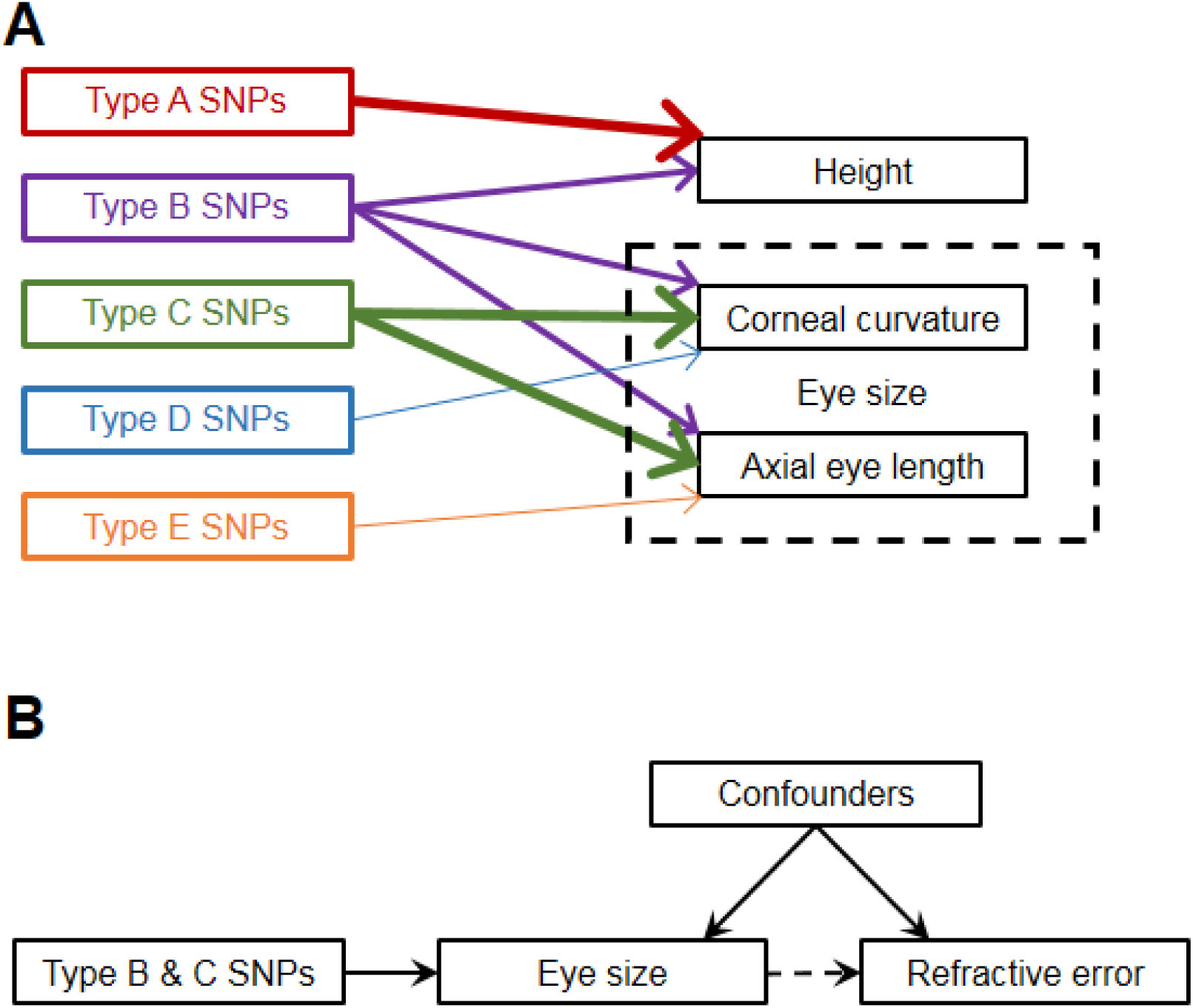
Causal diagrams (directed acyclic graphs). **Panel A**: examples of classes of genetic variant that exert an influence on height and/or eye size in emmetropes. Arrow thickness relates to variance explained by the class based on genetic correlations (e.g. in emmetropes the genetic correlation between corneal curvature and height « 0.30, while the genetic correlation between corneal curvature and axial length ≈ 0.85 [39]). Note that few genetic variants influence corneal curvature yet not axial length, and vice versa, i.e. most SNPs controlling axial length are 'Type C' SNPs, followed by 'Type B' SNPs. **Panel B**: Relationship between variables in Mendelian randomisation analysis; 'Type B & C' SNPs are used as instrumental variables to test for a causal relationship between eye size in emmetropes and refractive error.

